# Phase Gradients and Anisotropy of the Suprachiasmatic Network

**DOI:** 10.1101/2021.02.01.429173

**Authors:** Tomoko Yoshikawa, Scott Pauls, Nicholas Foley, Alana Taub, Joseph LeSauter, Duncan Foley, Ken-Ichi Honma, Sato Honma, Rae Silver

**Affiliations:** Organization for International Education and Exchange University of Toyama, Toyama 930-8555, Japan; Department of Mathematics, 6188 Kemeny Hall, Dartmouth College, Hanover, NH, 03755 USA; Department of Neuroscience and Zuckerman Mind Brain Behavior Institute, Columbia University, 3227 Broadway, New York City, NY, 10027, USA; Department of Psychology, Mail Code 5501, Columbia University, 1190 Amsterdam Avenue, New York City, NY, 10027, USA; Department of Neuroscience, Barnard College, New York City, NY, 10027, USA; Department of Economics, New School for Social Research, New York City, NY, 10011, USA; Research and Education Center for Brain Science, Hokkaido University, Sapporo, 060-8638, Japan; Center for Sleep and Circadian Rhythm Disorders, Sapporo Hanazono Hospital, Sapporo, 064-0915, Japan; Department of Neuroscience, Barnard College, 3009 Broadway, New York City, NY, 10027, USA; Department of Pathology and Cell Biology, Graduate teaching faculty, Columbia University Medical School, New York City, NY, 10032 USA

**Keywords:** suprachiasmatic nucleus, circadian, phase waves, network, connectome, vasopressin, vasoactive intestinal peptide, directionality, anisotropy, oscillation, phaseomes

## Abstract

Biological neural networks operate at several levels of granularity, from the individual neuron to local neural circuits to networks of thousands of cells. The daily oscillation of the brain’s master clock in the suprachiasmatic nucleus (SCN) rests on a yet to be identified network of connectivity among its ~20,000 neurons. The SCN provides an accessible model to explore neural organization at several levels of organization. To relate cellular to local and global network behaviors, we explore network topology by examining SCN slices in three orientations using immunochemistry, light and confocal microscopy, real-time imaging, and mathematical modeling. Importantly, the results reveal small local groupings of neurons that form intermediate structures, here termed “phaseomes” which can be identified through stable local phase differences of varying magnitude among neighboring cells. These local differences in phase are distinct from the global phase relationship – that between individual cells and the mean oscillation of the overall SCN. The magnitude of the phaseomes’ local phase differences are associated with a global phase gradient observed in the SCN’s rostral-caudal extent. Modeling results show that a gradient in connectivity strength can explain the observed gradient of phaseome strength, an extremely parsimonious explanation for the heterogeneous oscillatory structure of the SCN.

**Significance statement:** Oscillation is a fundamental property of information sensing and encoding in the brain. Using real time imaging and modeling, we explore encoding of time by examining circadian oscillation in single neurons, small groups of neurons, and the entire nucleus, in the brain’s master: the suprachiasmatic nucleus (SCN). New insights into the network organization underlying circadian rhythmicity include the discovery of intermediate structures, termed ‘phaseomes’, characterized by neurons which are stably out of phase with their neighbors. Modeling indicates that the pattern of phaseomes across the tissue encompasses a gradient in connectivity strength from the rostral to caudal aspects of the nucleus. Anisotropy in network organization emerges from comparisons of phaseomes and connectivity gradients in sagittal, horizontal and coronal slices.

## Introduction

It is widely accepted that the phasing of neuronal oscillation is an important aspect of network organization and brain function (1). The hypothalamic suprachiasmatic nuclei (SCN) function as a master circadian clock that orchestrates circadian rhythms in behavior and physiology. Each SCN is made up of ~10,000 neurons and the individual neurons contribute to circuits that support the coherent daily oscillation of the nucleus. While most SCN neurons express circadian oscillations, the individual cellular rhythms in the network are not synchronized in that they do not simultaneously reach peak phase (2–4). Orchestration of stable circadian rhythmicity requires a network that couples individual SCN neurons to each other (5–12)

With respect to circadian timing, a challenge is to understand how coherent daily rhythms emerge in the brain master clock through interactions of its individual neurons, ensembles of neurons, and larger-scale oscillation of the SCN tissue as a whole. Substantial evidence indicates that stable phase differences occur not only between adjacent neurons (13) but also among clusters of neurons in subregions of the nucleus (2, 14–20). While peak phase differs among neurons, relative phase does not drift (18), pointing to a non-uniform SCN network topology underlying tissue-wide oscillation.

Instead of synchronization of peak phase among individual elements, long-term, real-time luciferase reporter imaging of clock genes or proteins in SCN slices indicate phase waves that propagate over the entire nucleus with a ~24-hour rhythm. In coronal slices, these daily phase waves generally start in a distinct cluster of neurons in the arginine vasopressin-(AVP−) rich dorsal or dorsomedial region of the nucleus (2, 21). It is noteworthy that there are marked differences in oscillatory patterns among slices, likely due to inclusion of different network components included at the time of tissue harvesting. Within an individual slice however, the phase relationships of serial oscillatory waves are stable if the tissue is not perturbed (15). An important question is how these phase patterns link to the underlying fixed aspects of the SCN network. The localization of major clusters of SCN peptides do not fully explain the patterns of oscillation (2), and the precise topology of the SCN connectome has been difficult to establish in part because of the small size, dense packing and heterogeneity of its neurons and the fine caliber of fibers (22).

While understanding of the intra-SCN connectome is incomplete, the functional significance of the connections between two major regions, namely the ventral core and dorsal shell are well established (reviewed in (23)). The core-shell framework has produced both biological and modeling work that provides substantial insight into SCN oscillation (reviewed in (24)). An aspect of network topology that escapes notice in studies of core-shell relationships is the possibility that SCN networks are anisotropic and that key aspects of network topology are lost following transection of fibers that course rostro-caudally. Studies of other oscillatory networks, such as the thalamus, highlight the principle that network oscillatory properties differ when brain sections are cut in the transverse versus the longitudinal axes (25).

The goal of this study was to determine whether fixed properties of SCN tissue, specifically those set by the localization and connectivity of its neurons might underlie the observed SCN oscillatory phase patterns and their variations. How the intact SCN’s anatomy, morphology, connectivity gives rise to the phase relationships among SCN neurons or clusters of neurons remains elusive. Also elusive is how circadian oscillation is retained following ablation of major components of the nucleus (26, 27). To address some of the caveats in our understanding, we pair detailed morphological analyses of fixed tissue, studies of real time imaging of PER2::LUC expression in cultured SCN tissue and mathematical and statistical tools to explore SCN networks. We define the biological aspects of SCN organization that underlie the topography of individual cellular oscillations and investigate the impact of that evidence on simulations with a mathematical model. The biological results point to novel intermediate structures which we term ‘phaseomes’. Sagittal and horizontal slice orientations that maintain the SCN’s rostral-caudal axis reveal a global phase gradient associated with the magnitude of the local phase differences within the phaseomes. Modeling results show that a gradient of connectivity strengths between neurons can account for the observed phase gradient of the phaseomes along the rostral-caudal axis.

## Results

### Visualization of SCN in three planes

The SCN is made up of a heterogeneous population of neurons. To set the stage for understanding the relationship of regionally specific clusters of cell types to the SCN network topology, we first mapped the peptidergic organization of the mouse SCN, delineating the major peptidergic cell types that can be retained when SCN tissue is prepared in sagittal, horizontal or coronal orientations (Fig. 1A). The SCN is a bilateral structure, lying on each side of the 3^rd^ ventricle and extending approximately 350μm dorsoventrally, 300μm laterally from the third ventricle and 700-750μm rostrocaudally (including a finger-like rostral projection). The full rostro-caudal extent of the SCN is best seen in sagittal sections. The distribution of these key peptides aligns fully with the spatial distribution of corresponding genes (28). The peptide maps in Fig. 1 emphasize that the specific SCN neurons and networks captured when tissue is sectioned for ex vivo real time imaging of oscillation can differ markedly depending on the precise tilt of the brain when it is blocked and on the orientation in which it is sliced. These differences among slices provoke the question of which cellular and network components are necessary for oscillation and whether anisotropy (directionality) is a determinant of the pattern of SCN oscillation.

**Legend Fig. 1:**
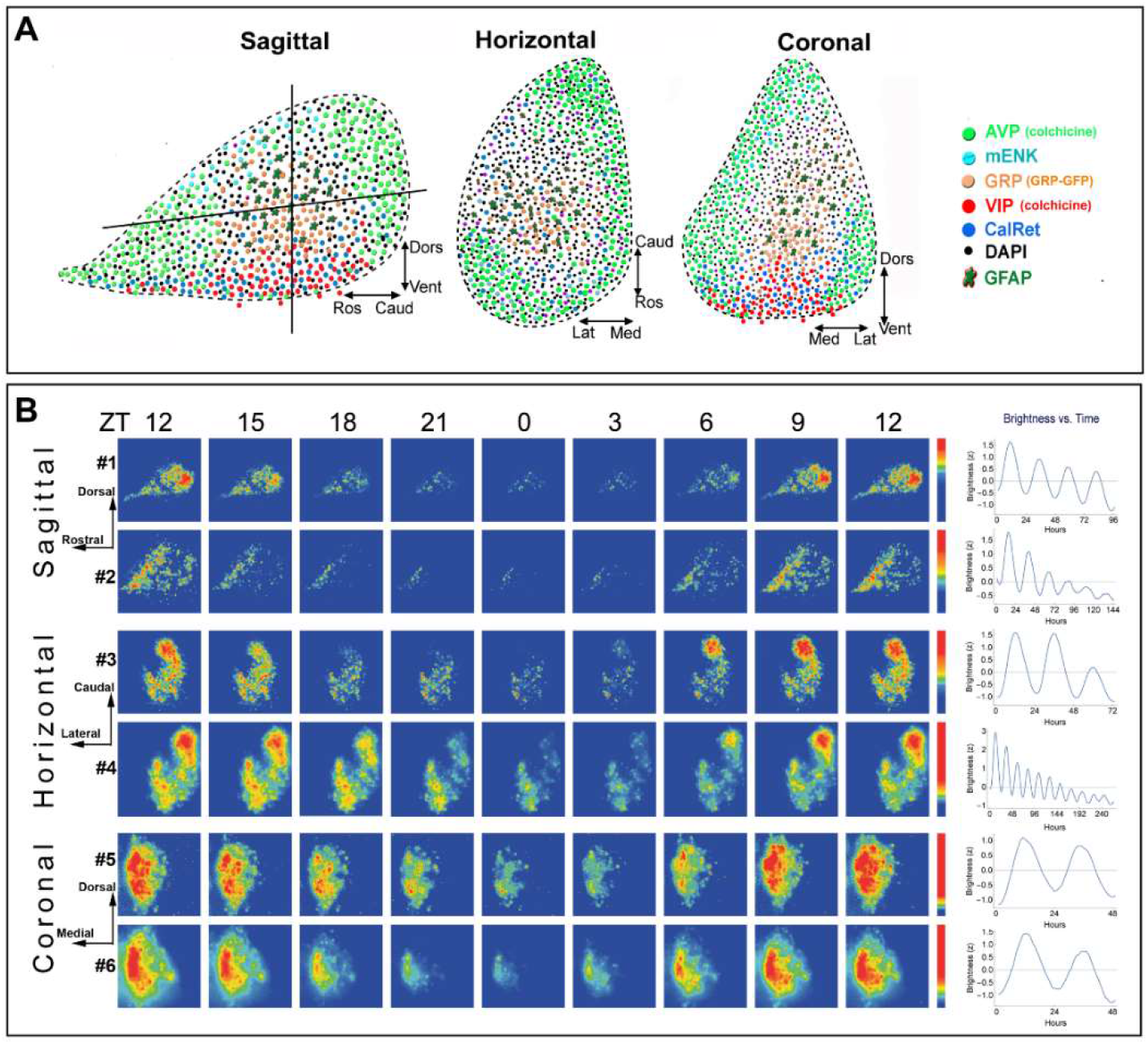
Architecture and PER2 expression of the SCN and Bioluminescence heat maps and brightness time series. **A.** Depiction of the peptidergic architecture by analysis of single-labelled SCN peptides in sagittal, coronal and horizontal planes. The localization of the major peptides of the SCN is shown in each orientation. The vertical and horizontal lines in the sagittal section indicates the plane shown in the adjacent cartoons of coronal and horizontal sections. Dapi label represents neurons that are not positive for any of the markers used. Vasoactive intestinal polypeptide (VIP)-containing neurons lie in the ventral core area. AVP neurons lie in the rostral protrusions and in much but not all of the outer borders of the nucleus. A gastrin-releasing peptide (GRP)-rich area, along with nearby glial fibrillary acidic protein-(GFAP-) positive elements, lies between the VIP core and AVP shell. In coronal sections the localization of core (VIP- and GRP-rich) and shell (AVP-rich) regions are seen. In horizontal sections, the precise peptidergic content of an SCN slice differs markedly depending on the angle and depth at which the SCN is cut; if the ventral aspect is included in a slice, then both the core and shell are represented. **B.** The spatiotemporal pattern in PER2∷LUC bioluminescence in SCN slices is shown at 3-h intervals for representative sagittal, coronal and horizontal slices. Time zero was defined as the time point with the lowest bioluminescent intensity (see Materials and Methods for details). The pseudocolored images are normalized to the brightest image of each slice. The rainbow scale (blue, low and red, high expression) for each slice lies on right side of the last panel. Mean circadian oscillation used in further analysis is shown in the right column. The SCN slices are numbered consistently so as to correspond on all figures.

### Effect of slice orientation on oscillation

We compared the effect of transecting SCN networks in three orientations by examining oscillation of PER2::LUC in sagittal, coronal and horizontal slices (Fig. 1B, middle panel). Imaging of sagittal sections has not previously been reported so we first examined whether the results seen in these slices correspond to data on PER2 expression in immunochemically stained sagittal sections harvested from animals sacrificed at specific circadian time points (Fig. S1). The results are confirmatory: The peak and trough oscillations are separated by ~12 hrs, and the overall oscillation has a period of ~24 hrs (see pseudocolored images of changes over time and quantification of the brightness time series in Fig. 1B, left and right respectively). The oscillation of the caudal SCN is more marked than the rostral aspect, and bears a different phase. The results for oscillation in our coronal and horizontal sections are consistent with previous reports on real time imaging of SCN slices (2, 20).

For real time imaging of the sagittal slices, we next investigated whether the anatomy of the slice, determined after imaging, impacted the production of oscillation or whether all slices oscillated irrespective of which circuit elements were present in the tissue. The results indicate that slices bearing both AVP and VIP neurons had robust rhythmicity while those lacking these peptides were not rhythmic (Fig. S2, see methods for oscillation criteria). Slices bearing a large number of AVP neurons but lacking VIP were not rhythmic, consistent with previous reports in coronal slices (26, 29).

### Observing phaseomes using biology and mathematics

In the real time imaging studies, close examination of the tissue near the trough of the oscillation reveals local phase heterogeneity with small populations of cells substantially out of phase with the surrounding tissue (Fig. 2A top panel). We designate these groupings as intermediate structures, here termed ‘phaseomes’. Phaseomes (denoted by red asterisks) are a local group of cells with stable phase heterogeneity, in which a cell is surrounded by a group of cells out of phase with it. A representative phaseome from the imaged material is shown at two time points with a phase difference of ~5 hours between the center cell and the surrounding cells (Fig. 2A middle and bottom panels). These phaseomes are not an artefact of the preparation, as they can also be seen in SCN sections from animals sacrificed near the trough of PER2 protein expression (Fig. S3).

**Legend Fig. 2:**
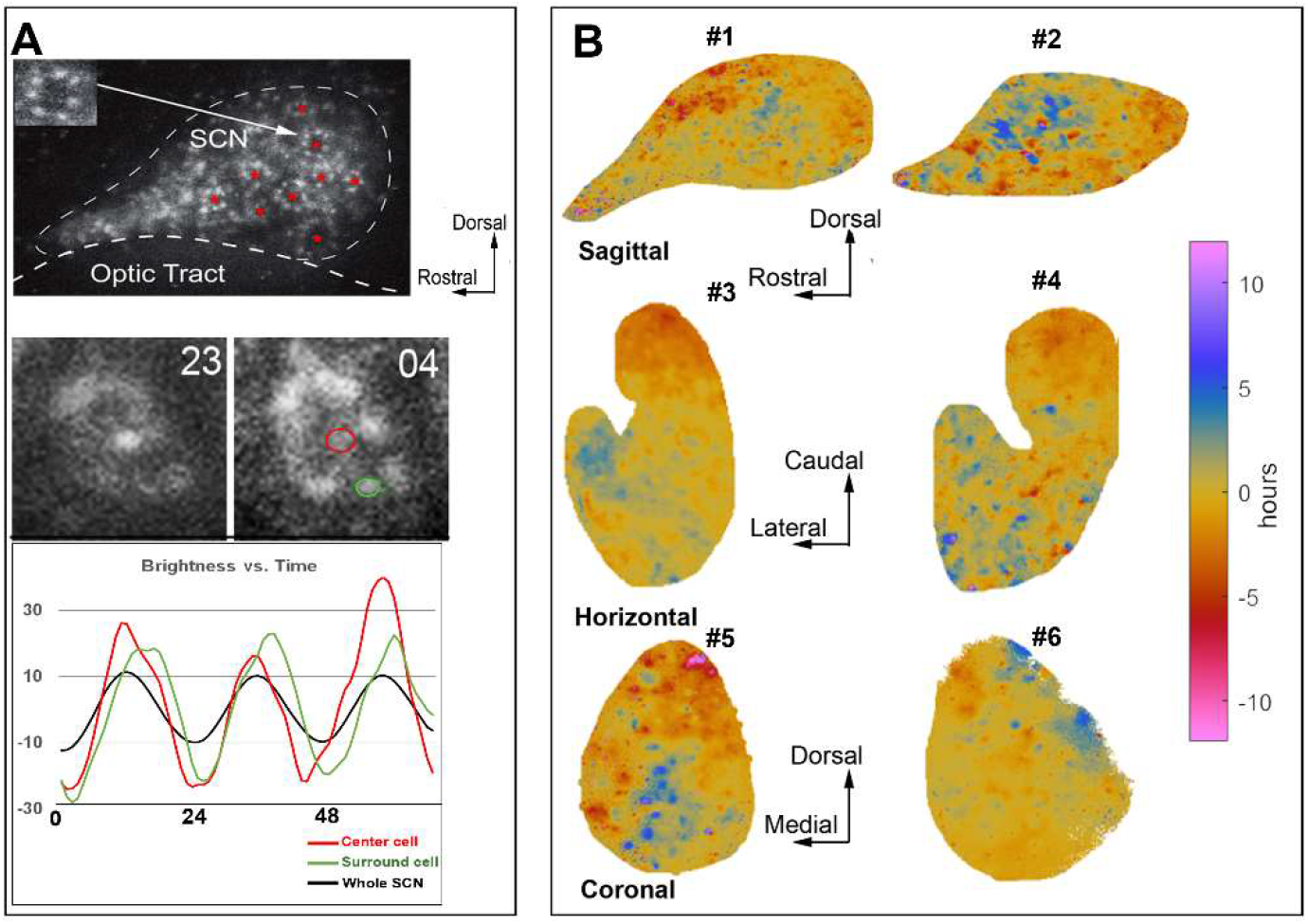
Global phase maps and phaseomes. **A. Phaseomes in the SCN.** The top panel is a raw image of a bioluminescent recording near the trough of PER2::LUC expression in a sagittal section. Red asterisks highlight the location of phaseomes. The inset shows a magnification of the phaseome indicated by the arrow. The middle panels are bioluminescent images of a phaseome in a sagittal section taken at 2 time points. At time 23, PER2::LUC expression is higher in the center cell compared to its neighbors. Five hours later, PER2::LUC expression is higher in surrounding cells compared to the center cell. The bottom panel shows the PER2::LUC oscillation over ~ 72hrs in the center cell (red circle),surround cell (green circle) and the whole SCN for the phaseome in the middle panel. **B.** Mathematically assessed **phase maps** for sagittal, horizontal and coronal slices of the SCN. Phase is represented by color, ranging from regions that are phase advanced (red) or phase delayed (blue) with respect to the tissue mean (yellow). The color of the 12 hour advanced regions matches to those with 12 hour delayed regions, as these will be in phase with one another in a 24 hr oscillation. All sections exhibit areas that are phase advanced and others that are phase delayed, often intermingled with one another.

### Relation of global to local phase

To explore global and local phase gradients through the full extent of the SCN, we devised a general analytical tool and applied it across different slice orientations. For global phase gradients, the phase of each pixel was assessed against the mean phase of the tissue (the global phase). This allowed determination of the effects on oscillation of preserving limited aspects of the network. Analysis of the temporal pattern of PER2::LUC expression in the slices through Fourier methods allowed identification of the phase of oscillation of any region of the SCN tissue relative to the mean circadian oscillation of the tissue as a whole (see Methods). Using such methods, we investigated whether phase maps differ by slice orientation and find that global phase maps show systematic phase anisotropy and heterogeneity (Fig. 2B).

Phase is represented by color, ranging from small regions that are phase advanced (red) or phase delayed (blue) with respect to the tissue mean (yellow). In the sagittal slices, there are neurons at the rostral pole and those in an area adjacent to the core that are phase delayed (blue speckled areas) relative to the mean tissue oscillation. Similarly, the horizontal sections also show near anti-phase relations between the rostral and caudal aspects of the tissue, with the caudal area substantially phase advanced (~5 hours) and the rostral aspect of the tissue substantially phase delayed (also ~5 hours) relative to the mean. In coronal sections we find, consistent with previous literature, a phase advanced region in the dorsal-medial aspect of the tissue. The rest of the phase map varies among slices [as previously reported (15, 17)], likely due to heterogenous sampling of the circuit depending on which part of the rostral-caudal extent to the slice was studied.

Phaseomes have not been previously reported but are consistent with prior work showing that adjacent neurons can be out of phase with each other (10). Local phase behavior may be a common occurrence across the SCN but could differ in structure depending on slice orientations and may be obscured by slice heterogeneity. The large global phase differences across the tissue confounds a local analysis – phase gaps of, for example, 4 hours might occur between adjacent regions with global phases at 10 hours and 6 hours from the mean in one part of the slice, and between regions with phases at 0 and −4 hours at another area. This consideration prompted us to develop a mathematical method to examine localized phase maps based on a center-surround filter that exposes relative local phase differences.

### Application of annular filter

A local phase analysis using a center-surround filter allows the examination of phaseomes computationally. This is illustrated by a very prominent phaseome identified from a global phase map in Fig. 2B that has a phase difference of about 10 hours. In a representative example the center cell size disk (blue) phase-lags the mean oscillation by about 5 hours (Fig. 3A top panel). Surrounding it are four neuron-sized regions (red), that lead the mean oscillation by about 5 hours. The mean phase of a cell-size disk of pixels was compared to that of a local annulus of equal radius around the disk (middle panel, see methods describing center-surround filter and “kernel”). This filter was then convolved over each pixel of the phase map, resulting in a local analysis (bottom panel). Phaseomes vary in the differences in the phase relationships of their components. The strength of the phaseomes vary across the tissue and we define the ‘strength’ of a phaseome as the magnitude of the phase gap between the center and surrounding cell-like regions. For example, a weak phaseome would have a center-surround phase gap of less than an hour in magnitude, while a strong one would be more than an hour (like the pictured example in Fig. 3A, bottom panel).

**Legend Fig. 3:**
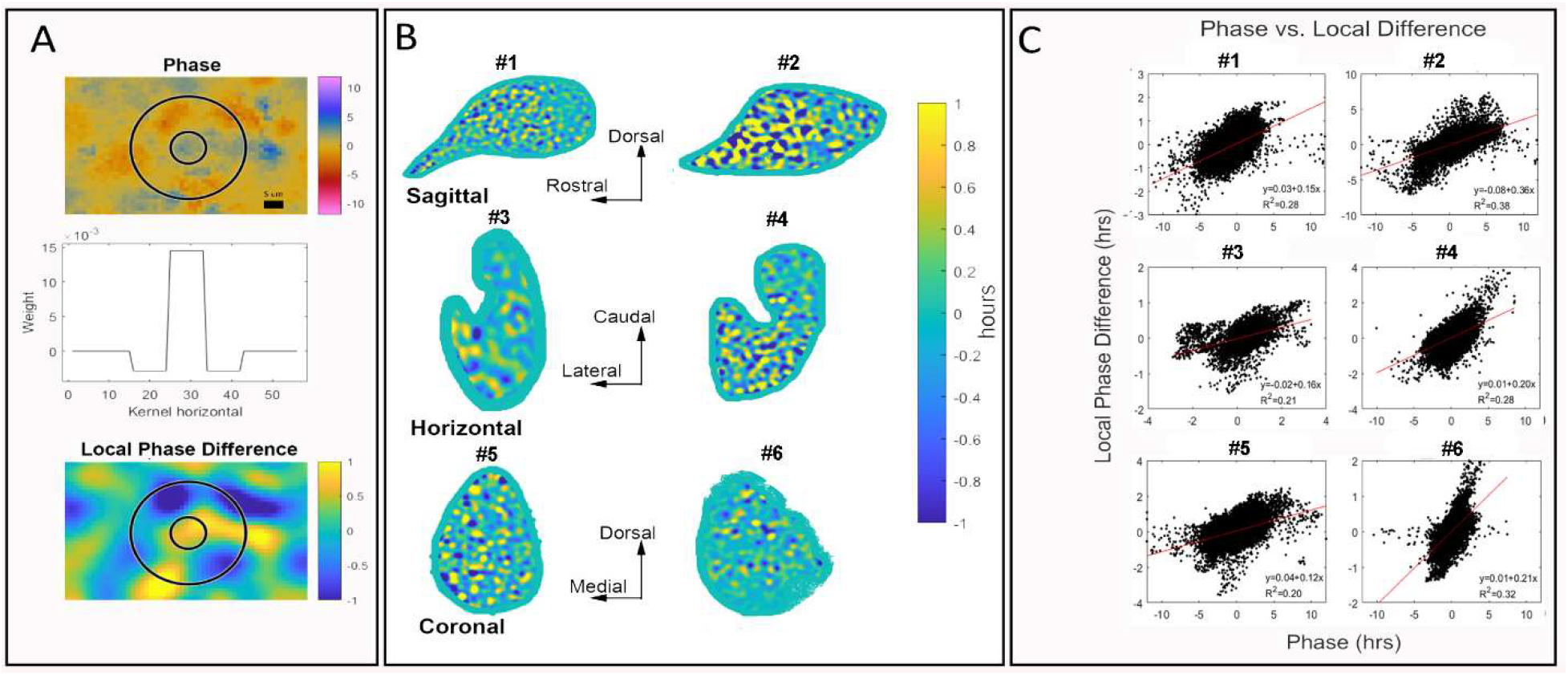
Local phase comparisons reveal phaseomes. **A.** Center-surround filter is superimposed on a magnified view of a phase map. Upper panel: In the center, there is a cell-sized region that is phase leading the mean signal. In the annular region several cell-sized areas are phase lagging the mean signal. This is consistent with evidence (Fig. 2) showing that neighboring areas have different levels of expression in similar local spatial arrangements. Middle panel: Cross-section of the center-surround filter used to compute the local phase differences. Lower panel: Results of the local phase difference computation on the same region and with the same center circle and surround annulus as the upper panel. The computation isolates the cell-sized regions and identifies them as phase leading or phase lagging the neighboring pixels, allowing a full slice analysis of local phase differences. **B.** Local phase differences as computed using the center-surround filter show numerous portions of the SCN that are out of phase with their neighboring tissue. Sagittal and horizontal slices show a gradient along the rostro-caudal axis where phase differences smaller at the caudal than the rostral aspect. The colors in this figure have been truncated to emphasize differences in values close to zero. The green border around the edge of each SCN is the result of “padding” to fill the kernel. **C.** Plot of the global phase against the local phase difference to examine the relationship between the two. Each point represents a single pixel. On the horizontal axis (global phase estimate of the tissue), positive values indicate oscillation that lags the mean oscillation while negative values indicate leading the mean oscillation. Linear regression lines (red) have high statistical significance and support the hypothesis that areas with larger local phase differences are more strongly leading or lagging the mean.

### Center-surround analyses of local phase comparisons reveal phaseomes throughout the SCN

Identifying the phaseomes computationally allowed us to determine whether local phase organization was reliant on the direction of the slice and/or the part of the tissue that was sampled over any extent (Fig. 3B). This is particularly relevant because intermediate structures have not been identified previously. The results indicate that phaseomes exist regardless of the orientation of the slices, but that their strength differs depending on the direction of slicing and the extent of the SCN examined, particularly on the rostral-caudal axis. The local areas have average phases that differ by roughly an hour regardless of slice orientation. This is shown in Fig. 3B, by the color in the convolved map which shows areas that are advanced (blue) and delayed (yellow) relative to the surrounding tissue mean (green). Note that phase differences in the local calculation are smaller than in the global calculation due to averaging. Many pixels in the global phase map (on which the kernel is convolved) are near the mean phase (i.e. phase difference of 0) and when included in the local kernel this brings the average closer to zero. We truncated the color scale to emphasize the pattern of local differences, even when the phaseomes are weak. The brightest yellow and darkest blue regions can have local differences larger than one or less than minus one respectively.

### The strength of phaseomes differs across the SCN’s rostral-caudal extent and among slice orientations

In the sagittal and horizontal sections phaseomes are stronger close to the rostral pole and weaker near the caudal extent (Fig. 3B, top two rows). We see this visually in the extent of the green areas (representing tissue mean) between the phase advanced (blue) and delayed (yellow) elements. In coronal sections, where the rostro-caudal extent is limited, we do not detect a consistent gradient on either the dorsal-ventral or medial-lateral axis (Fig. 3B, bottom row). As indicated in Fig 1A, the precise components of SCN tissue and the specific network components that are included in a slice will depend on precisely how it is blocked and cut. The coronal sections transect all rostro-caudal connections while this is not the case for sagittal and horizontal slices.

### The magnitude of local phase differences is related to the global phase deviation

Comparing the global phase estimates of the tissue (horizontal axis) to the local phase differences (vertical axis) reveals an interesting relationship (Fig. 3C): Areas that lead the mean oscillation by the largest amount tend to have large negative local phase differences, while those that lag by the largest amounts tend to have large positive local phase differences. Including the ordinary least squares regression line (in red) reveals positive linear slopes in each case. These results are statistically significant, with p-values estimated below machine tolerance in every case (all *p* < 10^−16^). If there were no relations between local phase differences and the global properties of the oscillation, we would expect no detectable correlation between global phase and local phase differences and the regression line would be horizontal. Instead, we see a positive correlation in each instance irrespective of orientation of the tissue.

### Kuramoto models connect the magnitude of local phase differences with the strength of local connectivity

We next asked what might be causing this rostral-caudal gradient and hypothesized that the topology of the SCN connectome that is retained when slicing in different directions gives rise to the observed gradients in phaseome strength. To test this hypothesis, we turn to mathematical modeling to examine the potential role of connectivity in the creation and strength of the local phase differences. We created a mathematical simulation encoding some of the properties of the SCN by constructing a 20 × 20 grid of oscillators with identical intrinsic frequencies that are linked to each of their four nearest neighbors. We change the connectivity by manipulating coupling strength, quadratically decreasing it as we move across the grid from right to left horizontally (Fig. 4A). In this depiction, the coupling strengths of two oscillators in the same vertical column are identical, but oscillators in the same row have different strengths depending on their locations in the grid.

**Legend Fig. 4:**
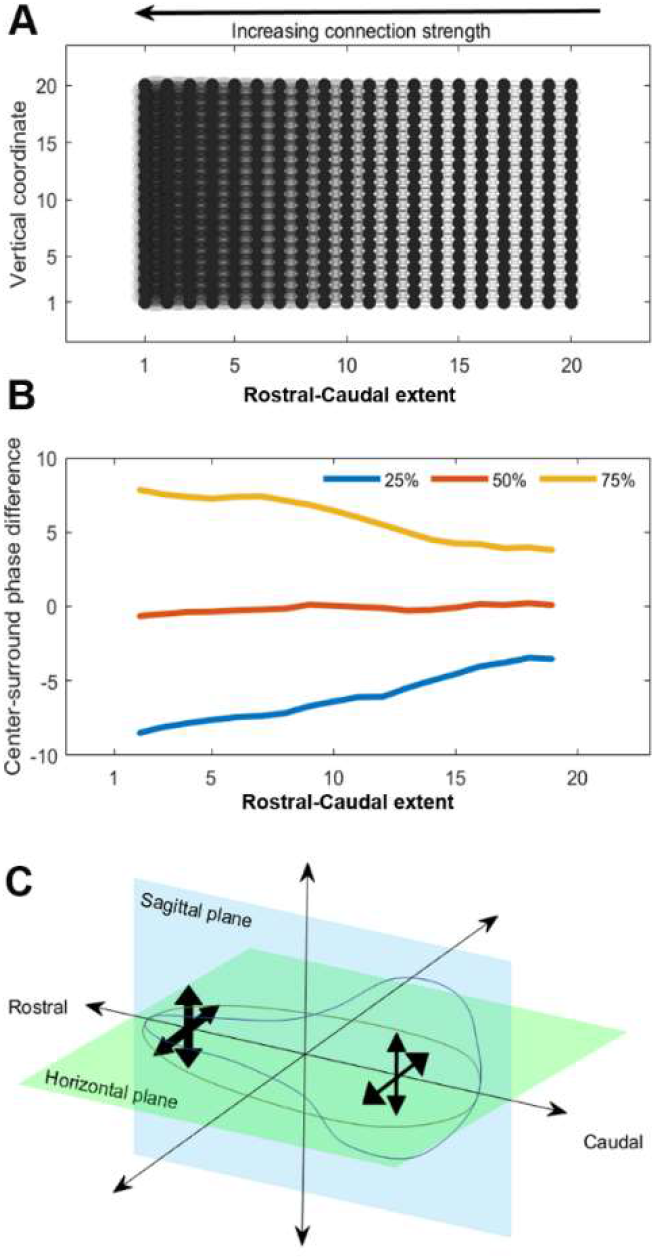
Kuramoto models connect the magnitude of local phase differences with the strength of local connectivity. **A.** A schematic of the network connectivity we use in simulations using the Kuramoto coupled oscillator model. The differences in shading from right to left indicate quadratically growing strength of the nearest neighbor coupling in the model. **B.** Results of applying the center-surround filter to simulated data over 500 trials. We report the quartiles of the distribution of local phase differences for each column of oscillators (as in A) as we move from right to left, showing that as the connectivity grows, so does the magnitude of the local phase differences. **C.** A schematic summarizing the interpretation of the local phase differences in the context of simulation results. Comparing to simulation results provides evidence that connection strength is weaker at the caudal edge of sagittal and horizontal sections and grows as we move towards the rostral tip.

We solved the associated Kuramoto system numerically over a period of 240 hours and calculated the phase of each oscillator (500 trials). We then applied a center-surround filter to the calculated phases, akin to the phaseome detector described above (Fig. 3A). For each vertical column of oscillators, we consider the distribution of phaseome strengths and compute their quartiles. Fig. 4B reports these quartiles as a function of the horizontal coordinate of the oscillator within the grid. We compare size of the difference between the 75^th^ percentile curve (yellow) and the 25^th^ percentile curve (blue) to assess the strength of the phaseomes as a function of coupling strength. The difference between the lower and upper quartiles is more compact on the right-hand than the left-hand side, indicating that the strength of the phaseomes decreases along with the strength of coupling. These results provide evidence for the hypothesis that the greater the phaseome strength, the stronger the local coupling between the oscillatory neurons (Fig. 4C).

## Discussion

### SCN networks as accessible models of oscillation

It is increasingly clear that systems in the brain responsible for temporal representation at many timescales rely on specific network organizations to sustain their activity (30). Network oscillations can bias input selection, temporally link neurons into dynamic assemblies, and modulate synaptic plasticity. The SCN is a uniquely accessible empirical model to study oscillatory networks: it is self-contained, it controls behavior and is reflected in observable physiological responses throughout the body.

### What is new in this work

The present work further demonstrates that while individual neurons oscillate, the rhythm in the SCN relies on the specific elements that are present in the network as a whole: the tissue is the issue. The observed oscillation in real time imaging of slices depends on what parts of the network are present after physical transection, and on the spatial and temporal resolution at which the tissue is being studied. Our use of biological, analytic and simulation tools demonstrate processes at multiple levels of analysis from individual cells to local and global organization in SCN networks and reveal the phaseome as an intermediate, local unit of organization. While previous research has not identified local phaseome-like structures in the SCN, the present findings are consistent with other locally identified neural units in the brain, such as the hypercolumns in area V1 (31) and striosomes of the striatum (32) that constitute intermediate structures parallel to the phaseome.

We have defined the strength of the phaseome as the magnitude of phase difference between the center and surrounding annulus of phase-locked cells. We find that the strength of phaseomes varies in a systematic gradient across the tissue if the rostral-caudal axis is preserved. Interestingly, this gradient of local organization of phaseomes is aligned with a previously reported global phase gradient along the same axis in tissue harvested from animals held in long daylengths (33). The overall strength of the phaseomes is greater in the rostral aspect, which tends to phase lag, and smaller in the caudal aspect which tends to phase lead in sagittal and horizontal slices (Fig. 2B, 3B).

Furthermore, strong phaseomes with negative local differences are associated with leading the global phase while strong phaseomes with positive differences lag it (Fig. 3C) – which is to say the relationship between local phase differences and global phase is positive regardless of how the tissue is cut. The linkage between local and global organization highlights the potential functional role of the phaseomes in integrating local oscillatory information – phaseomes are not merely physical structures, but a mesoscale component of the machinery that allows the SCN to construct and maintain a robust and consultable circadian rhythm.

Our consideration of phaseomes, by the nature of the data, is necessarily two-dimensional but in the full three-dimensional structure of the SCN the phaseomes may have more complex structures. While the phaseomes we observe in the PER2::LUC imaging movie frames appear as rosettes, their three-dimensional structures may take many possible topological types – spheres, cylinders, tori (34). Our methodology using the center-surround filter to help detect phaseomes privilege’s circularly phaseome, but the method can readily be adapted to explore other configurations by altering the filter. We propose an impressively parsimonious model for the cause of these local/global patterns seen in the rostro-caudal gradient of phaseome strength. The Kuramoto simulation results suggest that changes in strength of local coupling can produce similar patterns in model systems. Stronger local connectivity leads to stronger phaseomes in coupled oscillators (Fig. 4). It has been suggested that the brain clock network bears properties of small world networks (35–37), which have tight local coupling alongside some longer-range connectivity. Our work helps to delineate possible structures for the local coupling, connecting it functionally to properties of the oscillation across the tissue.

### Relationship to prior work

The results are consistent with previous descriptions of phase in clusters of SCN neurons. The occurrence of individual neurons having elevated PER2 protein at the overall trough of expression have previously been reported, though in prior work it was not known whether these cells oscillate in antiphase with the larger population, or whether they are arrhythmic and continually express PER (38–42). Our results indicate that an antiphase population is rhythmic, with high PER2 expression between ZT0 and ZT4 and low expression by ZT6 (Fig. S1).

While phase dispersal and phase waves have been described, the occurrence of phaseomes or other intermediate structures has not been noted previously. Perhaps these were seen but not investigated or, alternatively, this may be due to slice thickness. Our slices in the real time imaging preparations are thin (100μm), allowing for better cellular resolution, while in many other laboratories, slice thickness is ~300μm. With 300μm slices, it is difficult to visualize many individual cells simultaneously. Thin slices may have less of the global network than thick ones, but optimize single cell analysis. Another factor maybe the use of noise reduction algorithms or use of megapixels which also reduce the resolution. Choices within these algorithms include decisions on how many pixels to use to smooth the signal and how to smooth the signals which could have obscured phaseomes in prior reports.

### Importance of phase heterogeneity for timekeeping

Physiological and behavioral functions, including feeding, drinking, sleep-wake, body temperature, hormonal rhythms, and enzyme activity, have circadian rhythms with specific circadian peaks. To accurately assess circadian time at every time of day requires consulting cells whose PER2 concentrations are changing over time. Phase heterogeneity in PER2 expression allows this precise consultation throughout the circadian cycle, because at any phase of the mean oscillation, some cells will have swiftly changing expression of PER2. While the present work focuses on expression of PER2, the principle of heterogeneity in cellular rhythms applies more generally to cellular activity of a variety of responses. The precise consultation throughout the circadian cycle is enabled because at any phase of the mean oscillation, some cells will have rhythms at a particular phase. Our findings imply, in addition to sequentially phased PER2 rhythms, phaseomes may enable even more precise and specific regulations in overt rhythms.

An analogous finding in the visual system is that the neurons that provide the most information about the orientation of an edge are those whose firing rates change the most, rather than those that fire the most when presented with similarly oriented direction of motion (43). Protein concentrations of the mean signal in the SCN overall change slowly, especially when they are near the peak or trough of expression. Knowing the mean expression level of PER2 gives only a rough time signal – whether it is near the peak, trough or in between. Higher precision requires information complementary to mean concentration: the rate of change in concentrations of cells out of phase with the mean. Rapid changes in concentration within phasically heterogeneous cells provide continuously accurate time of day information regardless of the state of the mean oscillation. The advantages of this heterogeneity likely represent a general property of information encoding in the brain.

## Acknowledgments

The research was funded in part by grants from NSF | BIO | Division of Integrative Organismal Systems (IOS) 1256105; 1749500 (to RS and SP) and by JSPS KAKENHI (Grant 19K06774 to TY). TY thanks Dr. H. Gainer (National Institutes of Health) for the generous donation of AVP antibody for immunohistochemistry and Dr. Y. Shigeyoshi (Kindai University) for helpful discussions.

## Author Contributions

TY SPJLS AT conducted the experiments; TY RS SP DF JLS designed the experiments RS SP NF DF analyzed the data RS SP NF SH KH JLS DF wrote the paper.

## Methods

### Visualization of SCN peptides in sagittal, coronal and horizontal planes

To visualize the distribution of peptidergic cell types in the SCN, sections were stained for mENK, GRP, Calretinin, GFAP, dapi and AVP and VIP (the latter in colchicine treated mice) using the material and protocols previously reported in (44). To create the schematic, the localization of peptides and dapi was based on representative sections at the largest extent of the nucleus in each plane. For sagittal and coronal sections the localization of peptides and dapi was plotted based on a representative section at the largest extent of the nucleus, for the horizontal view, the data is based on previously unpublished material.

### Oscillation Criteria

As PER2::LUC expression in sagittal sections has previously never been described we asked whether circadian oscillation is seen in all slices harvested or alternatively, whether it is restricted to the slices that bore core and shell components. To this end, each slice was assessed to classify oscillation, independently by two observers. Also, slices were evaluated using Fourier analysis to determine statistical significance of the 24 hr period.

### SCN slice culture, bioluminescence

#### Slice culture

Mice were decapitated and enucleated after cervical dislocation between zeitgeber time (ZT) 5 and 9. The brain was removed and chilled in ice cold Hanks’ balanced salt solution, which was followed by serial slicing of the brain tissue (microslicer; Dosaka EM, Kyoto, Japan. Slices (100μm) were made on a sagittal, coronal or horizontal plane. The brain slices were cultured on a membrane (Millicell-CM membrane, Millipore) with 1.3 ml of DMEM containing 0.2 mM D-luciferin K and 5 % culture supplements as previously reported (33).

#### Bioluminescence recording

Images were obtained using a CCD camera cooled to −80 °C (ImagEM, Hamamatsu Photonics, Hamamatsu, Japan; iXon3, Andor, Belfast, UK). Bioluminescence was recorded hourly for 6 consecutive days. At the end of the recording period the brain slices were fixed with 4% paraformaldehyde in 0.1 M phosphate buffer (PFA) and prepared for immunohistochemistry.

### Visualization and analysis of luciferase in serial frames of the image stacks

The raw data consists of sequential images recording PER2::LUC expression over one hour intervals. Data processing includes restricting the images to a region of interest thereby delineating the SCN and windowing the time series to a range in which movement of the tissue is minimal. Further, image sequences were restricted from the first frame without movement to the longest possible multiple of 24 hours to alleviate artifacts in Fourier analyses. Raw images were imported into ImageJ (version 1.52). For visualization purposes, outliers were removed using manual observation of the histogram. The image was then imported into Photoshop, converted into RGB scale and a color gradient was applied. Next, the first peak of PER2::LUC expression time and each subsequent 3 hr interval was captured for a total of a 27 hr cycle. The brightness time series was computed as follows: 1) the data was restricted to the spatial ROI delineating the SCN, and the temporal ROI with minimal tissue movement. 2) The whole of the remaining data was then z-scored and plotted with the mean of each frame.

Only those with a robust 24 hr period were considered for further analysis. A further screening was then done using mean brightness time series, to ensure that there were several complete circadian oscillations in each slice. All slices meeting these criteria were further analyzed. We focused on sections that contained the rostral and caudal poles, as these preserved the full rostro-caudal extent of the nucleus, and included the rostral and caudal poles that cannot be studied in coronal sections. Two slices were chosen in each orientation to illustrate the results.

### Global scale phase map: Extracting the phase associated to the component of the signal with a 24-hour period (Figure 2B)

For each pixel, we computed a discrete Fourier transform of the time series (described above as a multiple of 24 hours). This results in each pixel having a complex number, *α* + *i β*, associated to the component of signal with a 24-hour period which allows us to compute the phase:

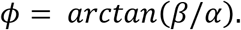

Each phase is given in radians, which we can convert to hours: 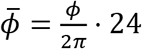. As phase is a relative statistic – it can only be measured against a baseline – we normalize the phases across the SCN so that a phase of zero corresponds with the mean signal across the SCN at the period of 24 hours. This results in the phase of every pixel being at most 12 hours phase advanced, or 12 hours phase delayed relative to the mean oscillation. The product of this process is a matrix *P* that aligns with the images in the frames of the PER2::LUC movies: *P*(*x*, *y*) is the phase extracted from the time series associated to the pixel in the (*x*, *y*) coordinate in a frame of the movie. Each lobe of the SCN was analyzed separately, and for visualization purposes one of the two lobes was chosen for each of the horizontal and coronal slices.

### Computing local scale phase difference with a center-surround filter

Examination of the global phase maps raises the question of the extent of heterogeneous phases in localized patches of the SCN. To focus on this local analysis, we compare the average of phases over a cell-sized disk of pixels to an average of those of an annular region surrounding that disk. This is done by convolving a filter isolating each putative cell-like region with the matrix of surrounding phase estimates. To this end, we use a binarized difference of Gaussians filter to facilitate the computation. Such a filter is also called a *center-surround* filter as it is positive on central disk and negative on a surround annulus.

We define the filter by

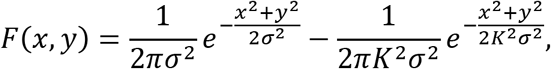

where (*x*, *y*) are the coordinates in the image plane, and *σ* and *K* are parameters that delineate the center and the surround: the first term is a Gaussian with standard deviation *σ*, defining the center, and the second a Gaussian with standard deviation *Kσ*, which defines the surround. For our purposes, using *σ* = 1, *K* = 2 creates a filter which is positive on a central disk of radius 4 pixels (roughly 9 um) and a surrounding annulus with outer radius 13 pixels (roughly 20 um) where the filter in negative. These sizes are consistent with comparing neurons to the surrounding tissue in our data, as SCN neuronal radii are 4-4.5 pixels. Figure 5A shows the shape of the center-surround filter superimposed on the phase maps of the tissue. Further, we binarize and normalize this filter by first replacing all values with absolute value less than 10^−5^ with zero. Then we replace the remaining positive values with +1 and negative values with −1. Last, we normalize the negative values of the filter by dividing by the number of pixels with a negative value and the positive values similarly. We denote the resulting filter by 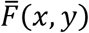. While we have chosen a circular annulus around a cell-like center based on patterns observed in the tissue (Fig. 4B,C), other surround kernels could be used to look for intermediate structures.

### Convolution of the center-surround kernel over the SCN

Convolving with this filter provides the difference in average phases between the disk and surrounding annulus centered at each pixel in the image. Applying center-surround filter to the SCN using 2D-convolution, 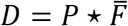, measures phase difference in the time series between each local disk and its nearest neighbors. The entry *D*(*x*, *y*) gives the difference between the average of the phases over the central disk of the filter, translated to be itself centered at the coordinates (*x*, *y*), and the average of the phases over the annular part of the filter.

### Kuramoto coupling model

We use Kuramoto model systems to investigate the possible contribution of connection strength to the existence of the rostral-caudal gradient in the local phase maps. Kuramoto systems with cluster synchronization, where smaller clusters of oscillators synchronize to different phases, exhibit higher intra-cluster and lower inter-cluster connectivity (45) suggesting similar features might hold for the clusters of tissue we observe in the local phase maps. The Kuramoto model systems comprise a set of oscillators that are connected to and influence one another.

Each oscillator in the system represents a neuron and is characterized by its intrinsic frequency, ω_*i*_, and the strength of its connections to other oscillators, {*a*_*i*1_, *a*_*i*2_, … , *a*_*in*_}. We represent the model system by a set of *n* differential equations:

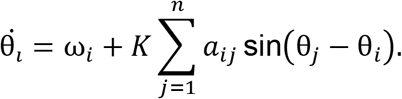

Here, *K* is a global underlying coupling strength. Using the Runge-Kutta method, we can numerically solve this set of equations for the {θ_*i*_}, allowing us to test different hypotheses. To look at the contribution of coupling to the rostral-caudal gradient, we set up a simple testing framework. First, oscillators are arranged on a grid and connected to their nearest neighbors. Second, we vary the strength of the connectivity in one direction to evaluate the effects of strength on the patterns of the resulting phases of the oscillators. To formalize this, we construct a model using a 20 × 20 grid of oscillators arranged on a planar lattice. We set the intrinsic frequencies to be the same, ω_*i*_ ≡ 2π/24, and *K* = 5. Letting (*x*(*i*), *y*(*i*)) be the planar coordinates of oscillator *i,* we define 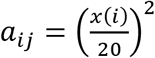 whenever *i* and *j* are neighboring oscillators in the plane (i.e. *max* { |*x*(*i*) − *x*(*j*)|, |*y*(*i*) − *y*(*j*)|}} = 1). This change in strength across the rectangle of oscillators models weaker connection strength on one side of the SCN that strengthens as we move across the tissue to the other side. Figure 6A shows a schematic of this setup. We solve numerically over a 240 hours period, in steps of 1 minute after providing initial conditions that are picked uniformly at random from [−π, π).

## Supplemental Material

## Supplementary Methods

### PER2 expression assessed by immunochemistry

#### Perfusion

Female (N = 11) and male (N = 25) mice were anesthetized with 100 mg/kg of ketamine and 10 mg/kg of xylazine 0 and sacrificed at 2 hour intervals throughout the 24h day. They were then perfused transcardially with 50 ml of saline followed by 75 ml of 4% paraformaldehyde. Brains were postfixed overnight at 4°C and then cryoprotected in 20% sucrose in 0.1 PB.

### Histology

Sections cut in the sagittal plane on a cryostat (Microm HM 500M, Walldorf, Germany) were collected into 0.1 M PBS. The brain sections were processed as free-floating sections in 24 wells plates. Sections were labeled for PER2 as follows: They were washed in 0.1M PB/0.1 % TX 100 and then blocked with 2% normal donkey serum (1:200; catalog #017-000-121 RRID: AB_2337258; Jackson ImmunoResearch) for 1 hr on a shaker at room temperature. They were placed in a 0.3% Triton-X 100 solution containing anti-PER2 antibody made in rabbit (1:500; catalog # AB2202; RRID:AB_1587380, EMD Millipore Corporation, Temecula, CA) at 4°C for 48 hrs. Sections were washed 3 times and incubated in a solution containing 0.3% Triton-X 100 and Cy3 donkey anti-rabbit (1:200; catalog # 711-165-152, RRID: AB_2307443 Jackson ImmunoResearch). Following a 2-hour incubation at room temperature, sections were washed in 0.1M PB and mounted, dehydrated in ethanol, cleared in CitriSolve (Fisher Scientific) and coverslipped with Krystalon (EM Diagnostics, Gibbstown, NJ).

### Microscopy and image analysis

PER2 expression was captured using a Nikon Eclipse E800 (Morrell Instruments, Melville, NY) microscope (excitation wavelengths 560±40 nm) equipped with a Nikon DSQi2 camera (Morrell Instruments) and using the Nikon NIS-elements basic research imaging software (RRID:SCR_002776; Morrell Instruments). Images were imported into Photoshop (RRID:SCR_014199; Adobe Photoshop CS. Berkeley, CA), “Shadows” and “Highlights” were adjusted in the “Image-Adjustment-Levels” dialog box in Photoshop, by dragging the black and white “Input Levels” to the edge of the first group of pixels on either end of the histogram.

### Anatomy of the slices after Imaging

Sagittal (n = 22 slices from 6 mice) slices were fixed with 4% PFA after the bioluminescence recording, and immunohistochemically labeled with a cocktail of antibodies against AVP (AVP-NP, PS419) and VIP (Peptide Institute, 14110). Immunohistochemical staining was examined by fluorescent microscopy (BZ9000; Keyence, Osaka, Japan) as previously reported (46). Anatomical analysis of peptide expression was conducted independently and prior to classification of rhythmicity. In Fig. 3, the number of positive cell bodies were scored as follows: For VIP: >10 cells = ++, 5-9 cells = +, 1-4 cells = ±, no cells = −. For AVP: >20 cells = ++, 5-20 cells = +, 1-5 cells = ±, no cell = −.

## Supplementary Figures

**Legend Fig. S1:**
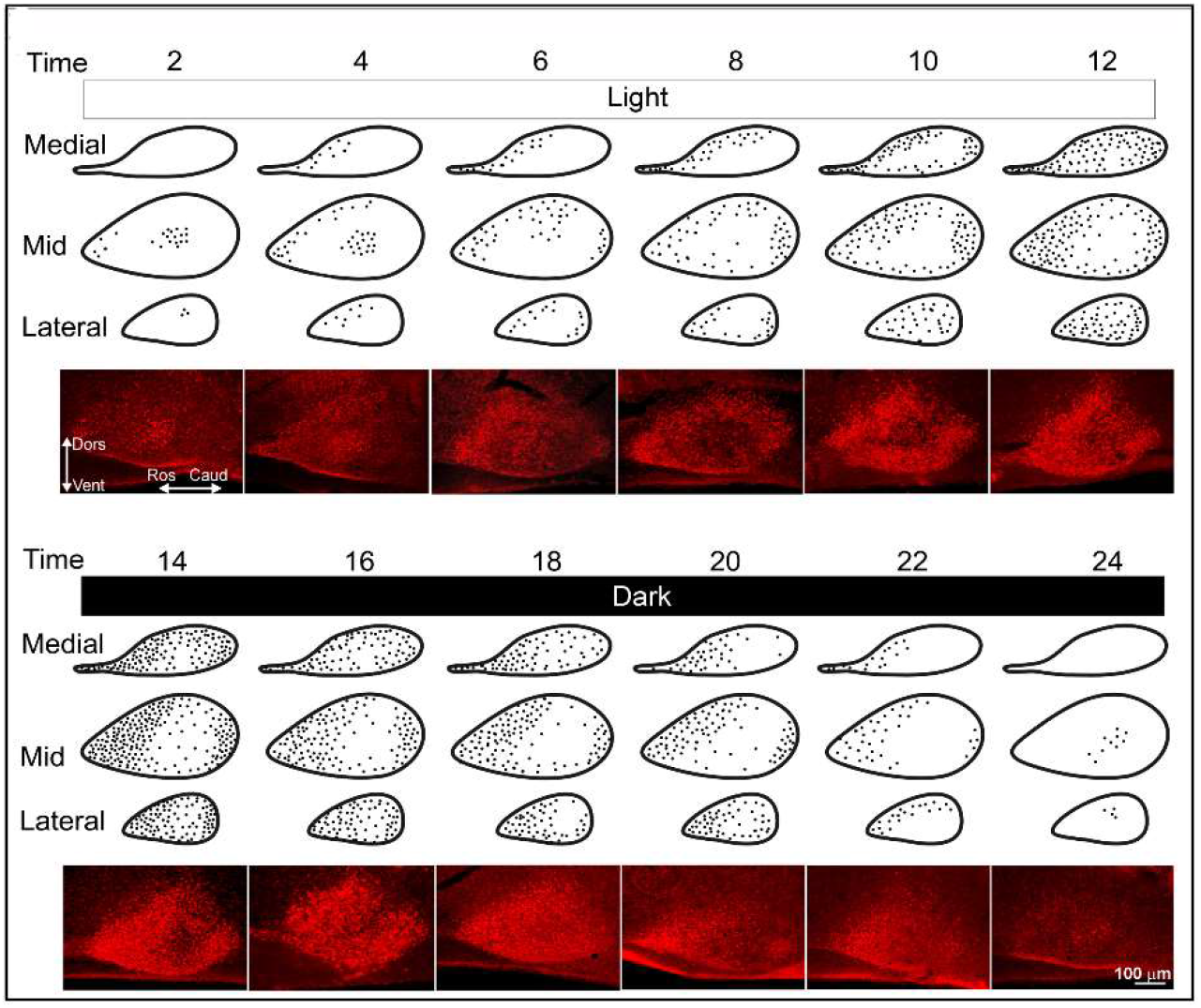
PER2 expression at 2 hr intervals in medial, mid- and lateral SCN. To localize changes in PER2 expression so as to have a baseline against which to compare the ex vivo sagittal slices in the real time imaging experiments, we assessed expression of the protein through the entire SCN in fixed tissue at 2 hr intervals. Sagittal sections of the SCN allowed visualization of the full rostro-caudal extent of the nucleus. Each dot represents the number and location of the PER2 nuclei observed in 2-3 brains at each time point. PER2-positive neurons can be seen at the trough of PER2 at ZT24/0 to ZT4 in the mid and lateral SCN in the mid-SCN. The rostral SCN expresses PER2 from ZT4 to ZT22. The photomicrographs in row 4 and 8 show the mid-SCN. The implication of regionally localized PER2 expression is that the observed network architectures will depend on the precise orientation of the slice. Maintaining the rostral and caudal poles of the SCN may preserve important circuit components that are lost in coronal sections.

**Legend Fig. S2:**
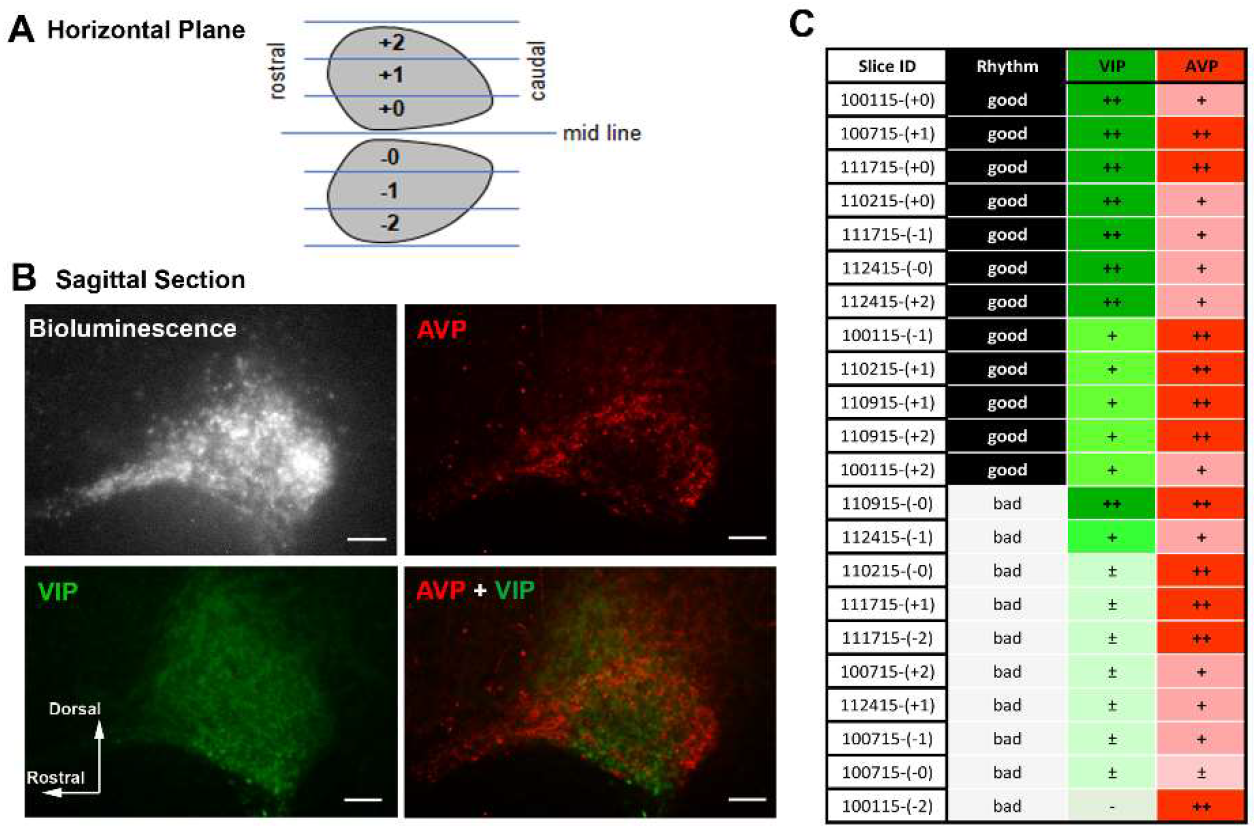
Post imaging immunohistochemical analysis of histology. **A.** Schematic drawing of the horizontal slice of the SCN. Blue lines are slicing plane of the sagittal slices. Numbers (−2 ~ +2) indicate position of the sagittal slices. **B.** Images of bioluminescence, immunohistochemical staining of AVP, VIP and overlay of AVP and VIP. **C.** Chart comparing robustness of oscillation and expression of AVP and VIP in each slice. Slice IDs indicate good (black) and poor (grey) rhythms. Number of immunopositive cell bodies for VIP and AVP are expressed in symbols. ++, high; +, medium; ±, low; −, none. Robust oscillation requires both AVP and VIP expression. Slices bearing a large number of AVP neurons but lacking VIP showed poor rhythm. The one slice that had both peptides could not classified with respect to rhythmicity due to technical equipment problems.

**Legend Fig. S3.**
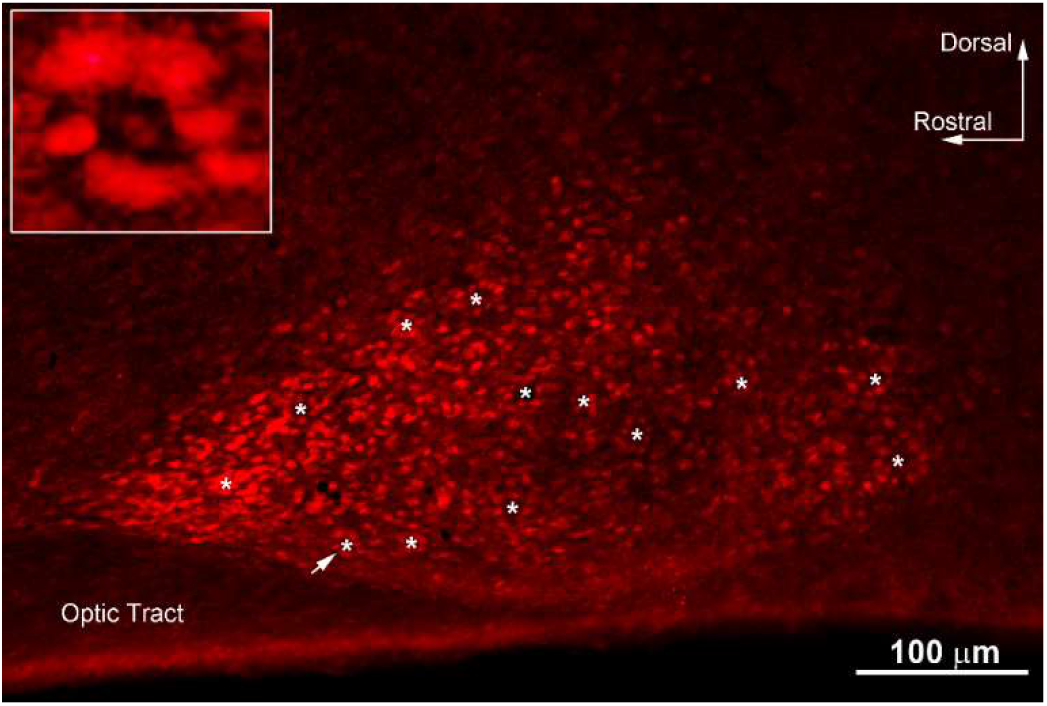
Phaseomes in fixed SCN tissue: Photomicrograph of a 50μm sagittal SCN section immunostained for PER2 (red) at ZT20. White asterisks show the location of phaseomes. The inset is a magnification of the phaseome indicated by the white arrow.

## References

1. G. Buzsaki, A. Draguhn, Neuronal oscillations in cortical networks. Science 304, 1926–1929 (2004).

2. J. A. Evans, T. L. Leise, O. Castanon-Cervantes, A. J. Davidson, Intrinsic Regulation of Spatiotemporal Organization within the Suprachiasmatic Nucleus. Plos One 6 (2011).

3. S. Koinuma et al., Regional circadian period difference in the suprachiasmatic nucleus of the mammalian circadian center. Eur J Neurosci 38, 2832–2841 (2013).

4. J. Schaap, C. M. Pennartz, J. H. Meijer, Electrophysiology of the circadian pacemaker in mammals. Chronobiology international 20, 171–188 (2003).

5. J. H. Abel et al., Functional network inference of the suprachiasmatic nucleus. Proceedings of the National Academy of Sciences 113, 4512–4517 (2016).

6. A. M. Finger, A. Kramer, Mammalian circadian systems: organization and modern life challenges. Acta Physiologica, e13548 (2020).

7. M. H. Hastings, E. S. Maywood, M. Brancaccio, Generation of circadian rhythms in the suprachiasmatic nucleus. Nature Reviews Neuroscience 19, 453–469 (2018).

8. S. Honma et al., “Suprachiasmatic nucleus: cellular clocks and networks” in Progress in brain research. (Elsevier, 2012), vol. 199, pp. 129–141.

9. I. T. Tokuda, D. Ono, S. Honma, K.-I. Honma, H. Herzel, Coherency of circadian rhythms in the SCN is governed by the interplay of two coupling factors. PLoS computational biology 14, e1006607 (2018).

10. S. Wang et al., Inferring dynamic topology for decoding spatiotemporal structures in complex heterogeneous networks. Proceedings of the National Academy of Sciences 115, 9300–9305 (2018).

11. A. B. Webb, N. Angelo, J. E. Huettner, E. D. Herzog, Intrinsic, nondeterministic circadian rhythm generation in identified mammalian neurons. P Natl Acad Sci USA 106, 16493–16498 (2009).

12. P. Indic, W. J. Schwartz, E. D. Herzog, N. C. Foley, M. C. Antle, Modeling the behavior of coupled cellular circadian oscillators in the suprachiasmatic nucleus. J Biol Rhythm 22, 211–219 (2007).

13. J. E. Quintero, S. J. Kuhlman, D. G. McMahon, The biological clock nucleus: A multiphasic oscillator network regulated by light. J Neurosci 23, 8070–8076 (2003).

14. T. M. Brown, H. D. Piggins, Spatiotemporal Heterogeneity in the Electrical Activity of Suprachiasmatic Nuclei Neurons and their Response to Photoperiod. J Biol Rhythm 24, 44–54 (2009).

15. N. C. Foley et al., Characterization of orderly spatiotemporal patterns of clock gene activation in mammalian suprachiasmatic nucleus. Eur J Neurosci 33, 1851–1865 (2011).

16. N. Inagaki, S. Honma, D. Ono, Y. Tanahashi, K. Honma, Separate oscillating cell groups in mouse suprachiasmatic nucleus couple photoperiodically to the onset and end of daily activity. P Natl Acad Sci USA 104, 7664–7669 (2007).

17. S. Pauls et al., Differential contributions of intra-cellular and inter-cellular mechanisms to the spatial and temporal architecture of the suprachiasmatic nucleus circadian circuitry in wild-type, cryptochrome-null and vasoactive intestinal peptide receptor 2-null mutant mice. Eur J Neurosci 40, 2528–2540 (2014).

18. S. Yamaguchi et al., Synchronization of cellular clocks in the suprachiasmatic nucleus. Science 302, 1408–1412 (2003).

19. L. L. Yan, H. Okamura, Gradients in the circadian expression of Per1 and Per2 genes in the rat suprachiasmatic nucleus. Eur J Neurosci 15, 1153–1162 (2002).

20. T. Yoshikawa et al., Localization of photoperiod responsive circadian oscillators in the mouse suprachiasmatic nucleus. Sci Rep-Uk 7 (2017).

21. R. Enoki et al., Topological specificity and hierarchical network of the circadian calcium rhythm in the suprachiasmatic nucleus. P Natl Acad Sci USA 109, 21498–21503 (2012).

22. A. N. Van den Pol, The Hypothalamic Suprachiasmatic Nucleus of Rat - Intrinsic Anatomy. J Comp Neurol 191, 661–702 (1980).

23. S. Honma, The mammalian circadian system: a hierarchical multi-oscillator structure for generating circadian rhythm. J Physiol Sci 68, 207–219 (2018).

24. S. D. Pauls, K. I. Honma, S. Honma, R. Silver, Deconstructing Circadian Rhythmicity with Models and Manipulations. Trends Neurosci 39, 405–419 (2016).

25. T. Gloveli et al., Orthogonal arrangement of rhythm-generating microcircuits in the hippocampus. P Natl Acad Sci USA 102, 13295–13300 (2005).

26. J. LeSauter, R. Silver, Localization of a suprachiasmatic nucleus subregion regulating locomotor rhythmicity. J Neurosci 19, 5574–5585 (1999).

27. B. Rusak, The role of the suprachiasmatic nuclei in the generation of circadian rhythms in the golden hamster, Mesocricetus auratus. Journal of comparative physiology 118, 145–164 (1977).

28. S. A. Wen et al., Spatiotemporal single-cell analysis of gene expression in the mouse suprachiasmatic nucleus. Nature Neuroscience 23, 456–+ (2020).

29. S. J. Aton, C. S. Colwell, A. J. Harmar, J. Waschek, E. D. Herzog, Vasoactive intestinal polypeptide mediates circadian rhythmicity and synchrony in mammalian clock neurons. Nature Neuroscience 8, 476–483 (2005).

30. G. Buzsâaki (2006) Rhythms of the brain. (Oxford University Press,, Oxford; New York).

31. D. H. Hubel, T. N. Wiesel, Receptive Fields and Functional Architecture of Monkey Striate Cortex. J Physiol-London 195, 215–& (1968).

32. A. M. Graybiel, C. W. Ragsdale, Histochemically distinct compartments in the striatum of human, monkeys, and cat demonstrated by acetylthiocholinesterase staining. Proceedings of the National Academy of Sciences 75, 5723–5726 (1978).

33. T. Yoshikawa et al., Daily exposure to cold phase-shifts the circadian clock of neonatal rats in vivo. Eur J Neurosci 37, 491–497 (2013).

34. M. J. Harding, H. F. McGraw, A. Nechiporuk, The roles and regulation of multicellular rosette structures during morphogenesis. Development 141, 2549–2558 (2014).

35. S. H. Hosseini, S. R. Kesler, Comparing connectivity pattern and small-world organization between structural correlation and resting-state networks in healthy adults. Neuroimage 78, 402–414 (2013).

36. S. H. Strogatz, Exploring complex networks. Nature 410, 268–276 (2001).

37. C. Vasalou, E. D. Herzog, M. A. Henson, Small-World Network Models of Intercellular Coupling Predict Enhanced Synchronization in the Suprachiasmatic Nucleus. J Biol Rhythm 24, 243–254 (2009).

38. M. D. Field et al., Analysis of clock proteins in mouse SCN demonstrates phylogenetic divergence of the circadian clockwork and resetting mechanisms. Neuron 25, 437–447 (2000).

39. M. H. Hastings, M. D. Field, E. S. Maywood, D. R. Weaver, S. M. Reppert, Differential Regulation of mPER1 and mTIM Proteins in the Mouse Suprachiasmatic Nuclei: New Insights into a Core Clock Mechanism. The Journal of Neuroscience 19, RC11–RC11 (1999).

40. E. D. Herzog, M. E. Geusz, S. B. S. Khalsa, M. Straume, G. D. Block, Circadian rhythms in mouse suprachiasmatic nucleus explants on multimicroelectrode plates. Brain research 757, 285–290 (1997).

41. J. LeSauter et al., A short half-life GFP mouse model for analysis of suprachiasmatic nucleus organization. Brain Res 964, 279–287 (2003).

42. K. Nakamura, A. Kitani, W. Strober, Cell contact–dependent immunosuppression by CD4+ CD25+ regulatory T cells is mediated by cell surface–bound transforming growth factor β. The Journal of experimental medicine 194, 629–644 (2001).

43. L. C. Osborne, W. Bialek, S. G. Lisberger, Time course of information about motion direction in visual area MT of macaque monkeys. J Neurosci 24, 3210–3222 (2004).

44. S. Varadarajan et al., Connectome of the Suprachiasmatic Nucleus: New Evidence of the Core-Shell Relationship. Eneuro 5 (2018).

45. C. Favaretto, A. Cenedese, F. Pasqualetti, Cluster Synchronization in Networks of Kuramoto Oscillators. Ifac Papersonline 50, 2433–2438 (2017).

46. T. Yoshikawa et al., Spatiotemporal profiles of arginine vasopressin transcription in cultured suprachiasmatic nucleus. Eur J Neurosci 42, 2678–2689 (2015).

